# SARS- CoV-2 viroporins: A multi-omics insight from nucleotides to amino acids

**DOI:** 10.1101/2021.09.26.461873

**Authors:** Manish Sarkar, Paul Etheimer, Victor Hannothiaux, Soham Saha

## Abstract

COVID-19 is caused by SARS-CoV-2 which has so far affected more than 500 million people worldwide and killed over 6 million as of 1st May, 2022. The approved emergency-use vaccines were lifesaving to such a devastating pandemic. Viroporins are important players of the life cycle of SARS-CoV-2 and are primary to its pathogenesis. We studied the two prominent viroporins of SARS-CoV-2 (i) Orf3a and (ii) Envelope (E) protein from a sequential and structural point of view. Orf3a is a cation selective viral ion channel which has been shown to disrupt the endosomal pathways. E protein is one of the most conserved proteins among the SARS-CoV proteome which affects the ERGIC related pathways. The aqueous medium through the viroporins mediates the non-selective translocation of cations, affecting ionic homeostasis in the host cellular compartments. This ionic imbalance could potentially lead to increased inflammatory response in the host cell. Our results shed light into the mechanism of viroporin action, which can be potentially leveraged for the development of antiviral therapeutics. Our results corroborate with previously published transcriptomic data from COVID-19 infected lung alveolar cells where inflammatory responses and molecular regulators directly impacted by ion channelling were upregulated.

## 1. Introduction

The COVID-19 (CoronaVIrus Disease 2019) is a severe acute respiratory syndrome (SARS) caused by a novel pathogenic β-coronaviral strain, SARS-CoV-2 which has affected millions all over the world. Vaccines have been developed using various technologies which have been approved by FDA like ChAdOx1 nCoV-19 (AstraZeneca) (1), mRNA-1273 (Moderna) (2), BNT162b2 (Pfizer) (3,4), Janssen Ad26.COV2.S (Janssen–Cilag International NV) (5), inactivated Vaccines (Vero Cell) (Sinopharm & Sinovac Life Sciences) (6,7) along with several other lead candidates in line. These vaccines are either whole inactivated viruses or have overlapping targets and origin which have been quite effective in mitigating the pandemic. SARS-CoV-2 has a genome encoding 28 proteins which play important roles in different stages of viral pathogenesis (8–11). The mRNA (Pfizer, Moderna), adenoviral vector-based (Astrazeneca, Janssen) and inactivated (Sinovac, Sinopharm) vaccines use epitopes from the spike protein to generate an immunogenic response in the body, thus creating an immunogenic memory. But since spike protein is very much prone to acquiring new mutations (12–14), these vaccines could lose their efficacies with the evolving viral genome (Center for Disease Control and Prevention). Emergence of numerous variants along with their sub-lineages of SARS-CoV-2 has been observed and is continuously being monitored (GISAID, W.H.O.). All of these variants have heavy mutational load on the spike protein (15–17), followed by the nucleocapsid (N) protein (15,18,19) and the RdRp (15,20–22). As a result, the cellular events involved in the viral pathogenesis, which are more conserved phylogenetically, become important. Ion channelling activity is one such feature which encompasses viroporins. These viroporins’ counter balance host cellular responses range from opposite directional ion flow to downstream disruptions of the host cell signalling pathways (23–25). The ion channelling activity of SARS-CoV-2 is prominently maintained by two proteins- (i) E protein (13,26) and (ii) Orf3a protein (26,27). The structure of the pentameric E protein from SARS-CoV-2 has been elucidated by solid state NMR (PDB id: 7K3G) (26) and homology modelling (13) which gives insights into its structure-function relationship and selectivity towards cations (28). Cryo-EM microscopy has given us insights into the dimeric structure of Orf3a (PDB id: 7KJR) which has two well defined aqueous channels fit for cation-selective channelling (29). Ion channelling mechanism of SARS-CoV-2 viroporins potentially leads to ionic imbalance and pH change of subcellular compartments of the infected host cells causing membrane disruption and intracellular malfunction. The E protein localises in the ERGIC membrane (27,30) while Orf3a localises primarily in the endosomal-lysosomal membranes of the human cellular host (31,32). Disruption of the ERGIC membrane affects protein translocation and processing of the host cell, causing ER stress (33) while endosomal-lysosomal membrane rupture leads to total breakdown of host trafficking machinery (23,32). Ionic imbalance and pH change (34) by the viroporins are important for the release of the virion particles as well.

Inflammatory responses in the host cell leading to ARDS could also be attributed to ionic imbalances (35). Mutations in pore forming regions of viroporins might lead to loss or gain in function of the ion channel partially or totally along the course of evolution of the viruses. However, the E protein and Orf3a exhibit a high degree of sequence identity between SARS-CoV-2 and its variants. We calculated the position specific frequency of all the residues of Orf3a and E protein using big data analysis to comment on the conservation of the sequences across the SARS-CoV-2 variants. The amino acid changes were identified, aligned and their homology models were structurally superimposed to each other. We inserted the viroporins (E and Orf3a) in organelle specific membrane mimics which allow solute and solvent movement, and performed molecular dynamics simulation studies to understand their mechanism of actions. While studying about the possible consequences, we found significant upregulation in immunomodulatory genes like CD40, IFNL1, IFNL2, IFNL3, IL12A, IL33, IL6 and NFkB1 after analysing the RNA sequencing dataset from Katsura et al., 2020 through our pipeline. Cellular functions like defence response to virus, regulation of cytokine production, response to IFN-γ and regulation of NFκB signalling were implicated in ontology analysis. Indeed, these cellular functions could be directly or indirectly affected by ionic imbalances in the cell, mediated by the viroporins Orf3a and E protein.

## 2. Materials and methods

### Position specific amino acid residue frequency calculation

Complete protein sequences were retrieved from NIH source data openly available (NCBI SARS-CoV-2 variant data packages; SARS-CoV-2 Resources - NCBI (nih.gov) and NCBI Virus; NCBI Virus (nih.gov) in the fasta format for different SARS CoV-2 variants. Sequences for the whole proteome from each of the variants were extracted, followed by separating the reading frames into Orf3a and E protein. Once the respective protein sequences for each variant were identified, we used the median length of the sequences as a selection filter. Any reported sequence lesser and greater than the median of the length of all proteins were discarded (incomplete or truncated sequences were hence excluded from the analysis). Indeed, for the proteins in each variant, we computed the following metric:

freq(Length protein < Median) * freq(Length protein > Median), and the results were always less than 10e-5.

This cut-off resulted in lower probability of having truncated and deleted sequences in our analysis. Using the refined list of amino acid sequences, we built a count matrix for each amino acid at each position for Orf3a and E protein for each variant. Using the count matrix, the positional frequency of each amino acid was computed and the amino acid with the highest frequency at each position was considered as the consensus sequence for the variant. The mutational frequency was calculated from the positional amino acid frequency using the following:

Mutational frequency = 1- positional frequency

The frequency in this calculation refers to the intra-variant frequency for each protein. All calculations were performed using custom made programs in Python. The data is represented as a heatmap across the different variants of SARS CoV-2 for the positional intra-variant frequency and as a stacked bar plot for the calculated mutational frequency for each amino acid position.

### Multiple sequence alignment

Consensus amino acid sequences of Orf3a and E protein were computed across the variants Q lineage of Alpha (Q.3), Beta (B.1.351), Gamma (P.1), Delta (B.1.617.2), AY lineage (AY.103) and Omicron (B.1.1.529). We aligned the variant specific mutation containing sequences with their references (**Orf3a**: NCBI Reference Sequence: YP_009724391.1; **E protein**: NCBI Reference Sequence: YP_009724392.1) in the online alignment tool Clustal Omega (62) and was visualised with Jalview (www.jalview.org).

### Homology modelling of the Orf3a protein and E protein

The structure of the Orf3a protein of SARS-CoV-2 has been elucidated using cryo-EM microscopy and determined as a dimeric protein (PDB id: 7KJR) (29). This structure has been used as a template for structure-based homology modelling of variant specific Orf3a structures. Pentameric E protein structure has been elucidated using solid state NMR from SARS-CoV-2 (PDB-id: 7K3G) (26) which has been templated to generate refined structural models for further analysis.

All the homology modelling procedures have been performed using MODELLER (64) and SWISS-MODEL (65). Model refinement and further structural fine-tuning of unreliable structural regions were done using the GalaxyWeB server (66). The structures obtained were validated by scores obtained from the MolProbity (67,68). The structures were chosen by comparing predominantly the different parameters like percentage of Ramachandran favoured and unfavoured residues, percentage of favoured and unfavoured rotamers, Mol-Probity score, and Clash Score, validating the quality of the modelled structures.

### Generation of the protein-membrane system

The transmembrane region of the proteins: Orf3a and Envelope (E) protein, was extracted and used for insertion in respective membrane mimics depending on their cellular localization. Orf3a from SARS-Cov-2 was inserted into an endosome mimicking membrane. The E protein was inserted into a membrane similar to the ERGIC. Asymmetric lipid compositions were maintained in the endosome and ERGIC mimicking systems. All the membrane insertion processes were performed in the CHARMM-GUI web server (69,70) similar to our previous study (13). The pore water of each of these channel proteins was removed while preparing the protein-membrane systems in CHARMM-GUI. The lipid compositions of each system corresponding to their inserted proteins are listed as follows:

### Molecular dynamics simulations using NAMD and VMD

The protein membrane systems were solvated using a 12 Å thick patch of TIP3P (71) waters at both sides of the protein bilayer complex along the *z* axis, and a uniform hexagonal area was maintained in the *x*–*y* plane. The K^+^ ion was added to the solvated system as required to mimic 0.15 M KCl which is similar to our physiological concentration of K^+^ ion. The structural models of proteins and lipids were presented using the CHARMM36 force field parameters (72,73) and NAMD 2.12 (74,75) was used to run the molecular dynamics (MD) simulations. Firstly, the energy of each system was minimised and then equilibrated using the NVT ensemble for 40 ps. The integration time step was kept at 1 fs with harmonic restraints of 10 kcal mol^−1^ Å^−2^ on the protein atoms and 5 kcal mol^−1^ Å^−2^ on the lipid headgroups. These are the first two steps of minimization and initial equilibration of the simulation system. Several cycles of NPT equilibration (four or more) were carried out after the first two steps with reducing force constants in each cycle to relax the restraints on the protein-membrane simulation system. The entire energy minimization and equilibration steps add up to around 2.25 ns for each simulation run. The minimised and equilibrated protein-membrane system was then simulated for 5 ns using an integrating time step of 2 fs, constraining all H-containing bonds by the SHAKE algorithm (76). The total sampling time of the trajectories altogether added to ∼20 ns. Langevin dynamics was used in all the simulations to keep the temperature constant at 303 K with a damping coefficient of 1 ps^−1^, and the Langevin piston method was used in NPT ensembles to keep the pressure constant at 1 atm with a coupling constant of τP = 0.5 ps (77). In all these simulations, short-range nonbonded interactions were switched off between 10 and 12 Å. The Particle Mesh Ewald method (78) was employed with a grid size of 1 Å for the estimation of long-range electrostatic interactions. The total energy of the simulation system, number of H-bonds, RMSD, RMSF and solvent accessible surface area (SASA) of specific pore forming residues were analysed with respect to time steps as obtained from the results of the NAMD simulation in VMD (79) interface and snapshots of the timesteps were represented and visualised using Chimera 1.10 (80).

### Transcriptomic analysis

Transcriptomic data was reanalyzed from the following source: *Katsura et al*., *2020 Cell Stem Cell* (36). The data is derived from whole genome RNA sequencing from a modular alveolo-sphere culture system of human alveolar type 2 cells/pneumocytes derived from primary lung tissue (36). Data was downloaded from Gene Expression Omnibus library [ID: GSE152586] and pre-processing of the fastq files was performed. Details of data extraction and experimental procedures are available in the original publication: *Katsura et al*., *2020 Cell Stem Cell*. The DESeq2 package in R BioConductor (http://www.bioconductor.org/packages/release/bioc/html/DESeq2.html) was used to analyse the data. The normalised data was used for visualisation, and differential analysis of the count data (81). The DESeq2 data class consists of a count matrix with rows corresponding to genes and columns denoting experimental samples (control and COVID-19). For dimensional reduction and outlier identification, we performed a principal component analysis (PCA) on the DESeq2 data class of count reads. The details of the *DESeq2* pipeline are discussed in detail in *Love et al*., *2014 Genome Biology* (81). Briefly, *DESeq2* package models the data counts on the count matrix using a gamma-Poisson distribution with mean (normalised concentration of cDNA fragments from the gene in a sample). The size factors are determined by the median-of-ratios method. For each gene, a generalised linear model (GLM), which returns overall expression strength of the gene, log_2_ of the fold change (LFC) between the two groups compared. The p values of comparison between control and infected samples are adjusted for multiple testing using the Benjamini and Hochberg procedure.

To identify the processes encoded by the upregulated genes, we used the publicly available protocol in Metascape (www.metascape.org/; (82)). We annotated the functions encoded by the genes using the following gene ontology enrichment: Biological processes, Cellular components, Molecular components, KEGG pathway and Reactome pathways. Metascape combines functional enrichment, interactome analysis, gene annotation, and membership search to leverage over 40 independent knowledge bases. The minimum overlap was kept at 3, the p value cut-off at 0.01 and the minimum enrichment was kept at 1.5. The network type was set at full network, network edges defined by confidence of the highest threshold (90%).

### Statistical analysis

For statistical comparisons, we have used Kolmogorov-Smirnov (KS) Test for comparing the cumulative distributions and unpaired students’ t-test for comparison between different conditions. The p-values reported in this paper consider p < 0.05 to be statistically significant.

## 3. Results

### 3.1. Intra-variant sequence analysis of SARS-CoV-2 viroporins, Orf3a and Envelope (E) protein

We have developed a new analytical program to identify invariant sequences conserved among the currently sequenced genomes of SARS CoV-2 in light of Orf3a and E protein (Figure 1A). We extracted the sequences of Orf3a and E proteins from the following variants respectively: Alpha (B.1.1.7; Figure 1B; N = 105187 and 105169), Q lineage of Alpha (Q.3; Figure 1B; N = 5300 and 5293) Beta (B.1.351; Figure 1B; N = 3814 and 3209), Gamma (P.1; Figure 1B; N = 13593 and 13558), Delta (B.1.617.2; Figure 1B; N = 50397 and 50198), AY lineage of Delta (AY.103; Figure 1B; N = 48918 in both the proteins), Omicron (B.1.1.529; Figure 1B; N = 468 in both the proteins) and BA lineage of Omicron (BA.1.1; Figure 1B; N = 167839 in both the proteins). Sequences greater than or equal to the median length were used for creating the count matrix (Figure 1B). The count matrix generated for Orf3a intra-variant position-specific frequency for each amino acid showed certain positions to have lower frequencies compared to a stable score of 1 (Figure 1C). This fluctuation in position specific frequencies was very low (range: 0-0.148), and hence could not be assigned as a stable mutation (Figure 1D). In order to ascertain the stability of mutations observed, we calculated the consensus sequences for Orf3a and E protein for each variant. Additionally, a similar count matrix generated for E protein intra-variant position-specific frequency also showed lower frequencies in certain positions compared to a stable score of 1 (Figure 1E). The fluctuation in position specific frequencies was quite low (range: 0-0.064), much lower than its Orf3a counterpart (Figure 1F).

**Figure 1.**
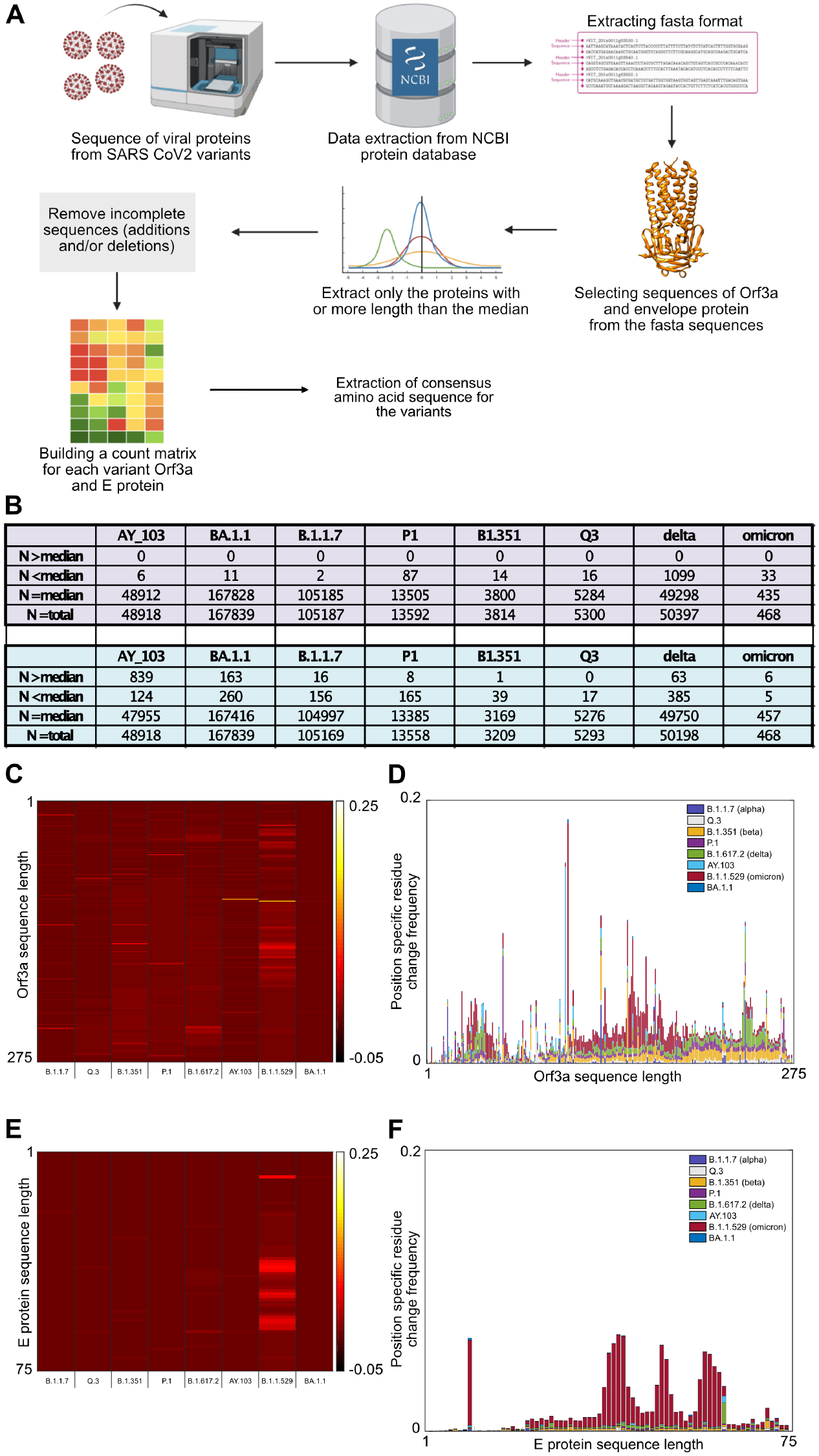
A. Analytical pipeline for determining the consensus sequences for Orf3a and E protein for the SARS-CoV-2 variants. The pipeline describes the steps from extraction of protein sequences for each protein to the determination of the position specific residue frequency. B. Tables showing the number of sequences excluded and included in the analysis using a median length threshold. C. Heatmap showing the position-specific residue intravariant frequency of Orf3q protein for each SARS-CoV-2 variant used in the analysis. Heatmap scale: - 0.05 to 0.25. D. A stacked bar plot showing the position-specific residue change frequency of Orf3q protein for each SARS-CoV-2 variant used in the analysis. E. Heatmap showing the position-specific residue intravariant frequency of E protein for each SARS-CoV-2 variant used in the analysis. Heatmap scale: - 0.05 to 0.25. F. A stacked bar plot showing the position-specific residue change frequency of E protein for each SARS-CoV-2 variant used in the analysis.

We computed the position-specific residue frequency and observed the position S26L of Orf3a in B.1.617.2 (Delta) and T9I of E protein in B.1.1.529 (Omicron) and BA.1.1 (BA lineage of Omicron), to have relatively higher scores (Figure 1D, F).

### 3.2. Structural modelling and analysis of representative mutations in Orf3a across SARS-CoV-2 variants

In correlation to previous results, we have identified representative mutations across Orf3a sequence specific for different SARS-CoV-2 variants (Figure 2A). All the variants except the Alpha, have one or more random mutations. The structure of Orf3a dimer has been elucidated using Cryo-EM at 2.1 angstrom, which spans from 40^th^ to 236^th^ amino acid, in complex with human apolipoprotein A (PDB id: 7KJR). The dimeric structure of Orf3a was extracted from the pdb file which was used as the reference (Figure 2B) for homology modelling of the representative Orf3a structures across SARS-CoV-2 variants (Supplementary figure 1). The refined Orf3a structures of the variants with the best parameters i.e.-Clash score, Molprobity score, Ramachandran outliers, Rotamer outliers, were chosen for further analysis of structural overlap. The Orf3a reference structure was superimposed on modelled structures from Alpha (B.1.1.7; Supplementary Figure 1A; RMSD = 0.89; TM score = 0.953), Q lineage of Alpha (Q.3; Supplementary Figure 1B; RMSD = 1.14 ; TM score = 0.945) Beta (B.1.351; Supplementary Figure 1C; RMSD = 1.19 ; TM score = 0.942), Gamma (P.1; Supplementary Figure 1D; RMSD = 0.84 ; TM score = 0.954), Delta (B.1.617.2; Supplementary Figure 1E; RMSD = 0 ; TM score = 1), AY lineage of Delta (AY.103; Supplementary Figure 1F; RMSD = 1.10 ; TM score = 0.945), Omicron (B.1.1.529; Supplementary Figure 1G; RMSD = 0.90 ; TM score = 0.952) and BA lineage of Omicron (BA.1.1; Supplementary Figure 1H; RMSD = 0.96 ; TM score = 0.951). The reference structures are shown in orange and the variants are shown in cyan (Supplementary Figure 1). According to our previous result, the Alpha and BA sub-lineage of Omicron do not have any significant mutations. In other variants, the following representative mutations were analysed in the Orf3a dimer overlap across variants: Beta-Q57H (Figure 2C-i; RMSD = 1.19; TM score = 0.942), S171L (Figure 2C-ii; RMSD = 1.19; TM score = 0.942); AY lineage of Delta-P104S (Figure 2C-iii; RMSD = 1.10 ; TM score = 0.945); Omicron-L106F (Figure 2C-iv; RMSD = 0.90 ; TM score = 0.952), S165F (Figure 2C-v; RMSD = 0.90 ; TM score = 0.952). However, the mutations present in the gamma (S253P) and delta variant (S26L) of Orf3a are in positions not included in the cryo-EM reference structure (40-236) and thus could not be mapped. The TM-score matrix indicates that there is no major structural variability in Orf3a protein among the variants (Figure 2D).

**Figure 2.**
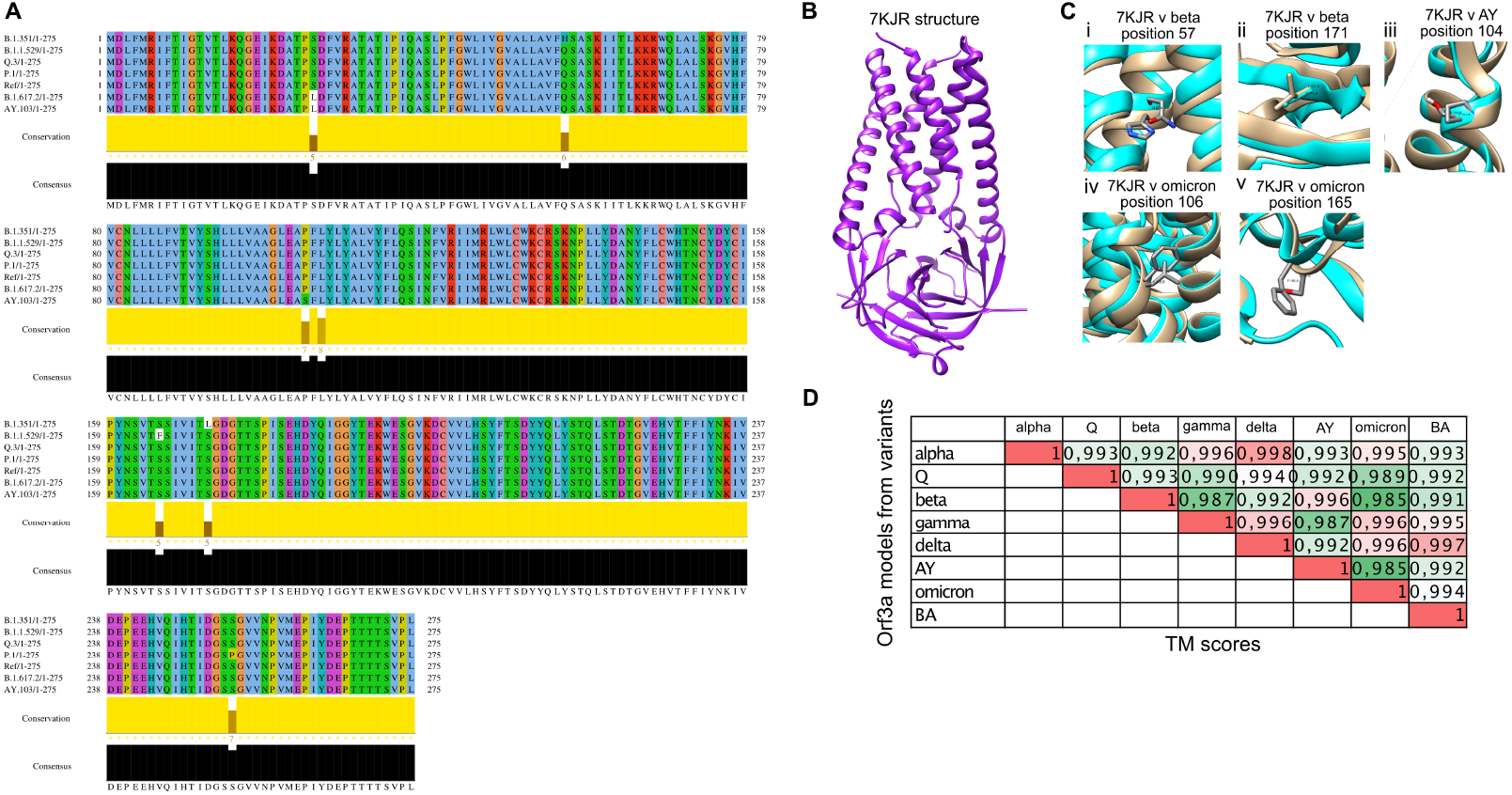
A. Multiple sequence alignment of representative mutations containing consensus sequences across all variants (except Alpha) of Orf3a protein. B. Reference structure: Cryo-EM structure of Orf3a protein dimer (PDB id: 7KJR). C. Structural superposition of variant specific mutations of Orf3a in SARS-CoV-2 variants with the reference structure: (i) Beta- Q57H (ii) Beta- S171L (iii) AY lineage of Delta- P104S (iv) Omicron- L106F (v) Omicron - S165F D. TM- score matrix of Orf3a protein across SARS-CoV-2 variants.

### 3.3. Molecular dynamics simulation of the TM region of Orf3a (40-125) from SARS-CoV-2 in endosomal membrane mimicking system

The SARS-CoV-2 Orf3a dimeric protein structure has been determined using cryo-EM at 2.0 angstroms resolution (PDB id: 7KJR) (29). The transmembrane region of the protein ranges from 40 to 125th residue of the protein. We performed membrane insertion of the truncated structure of Orf3a (40-125) using CHARMM-GUI. It comprises lipid components mimicking the human endosomal membrane (Table 1). Then a molecular dynamics simulation of 5 ns was performed using NAMD to understand the channelling activity of the upper and lower channels of the protein (Supplementary Movie 1). We observed the presence of the upper channel in the protein-membrane system (Figure 3A-black dotted circle). The water dynamics were observed at regular intervals of 0.5 ns for 2 ns from the initial timestep starting at 0 ns (Figure 3Bi), 0.5 ns (Figure 3Bii), 1 ns (Figure 3Biii), 1.5 ns (Figure 3Biv), and 2 ns (Figure 3Bv). Similarly, the lower channel (Figure 3C-black dotted circle) was also observed at similar timesteps (Figure 3i-v). We calculated the number of H-bonds (Figure 3E), RMSD (Figure 3F), total energy (Figure 3G), and RMSF (Figure 3H) for 5 ns of the simulation. The RMSD remained below 2.5 angstroms ((Figure 3F), indicating that the protein-membrane system has low structural variability. In addition, the total energy of the system remains largely unchanged at -5.7×10^4^ kcal/mol (Figure 3G) throughout the time period of the simulation.

**Table 1:**
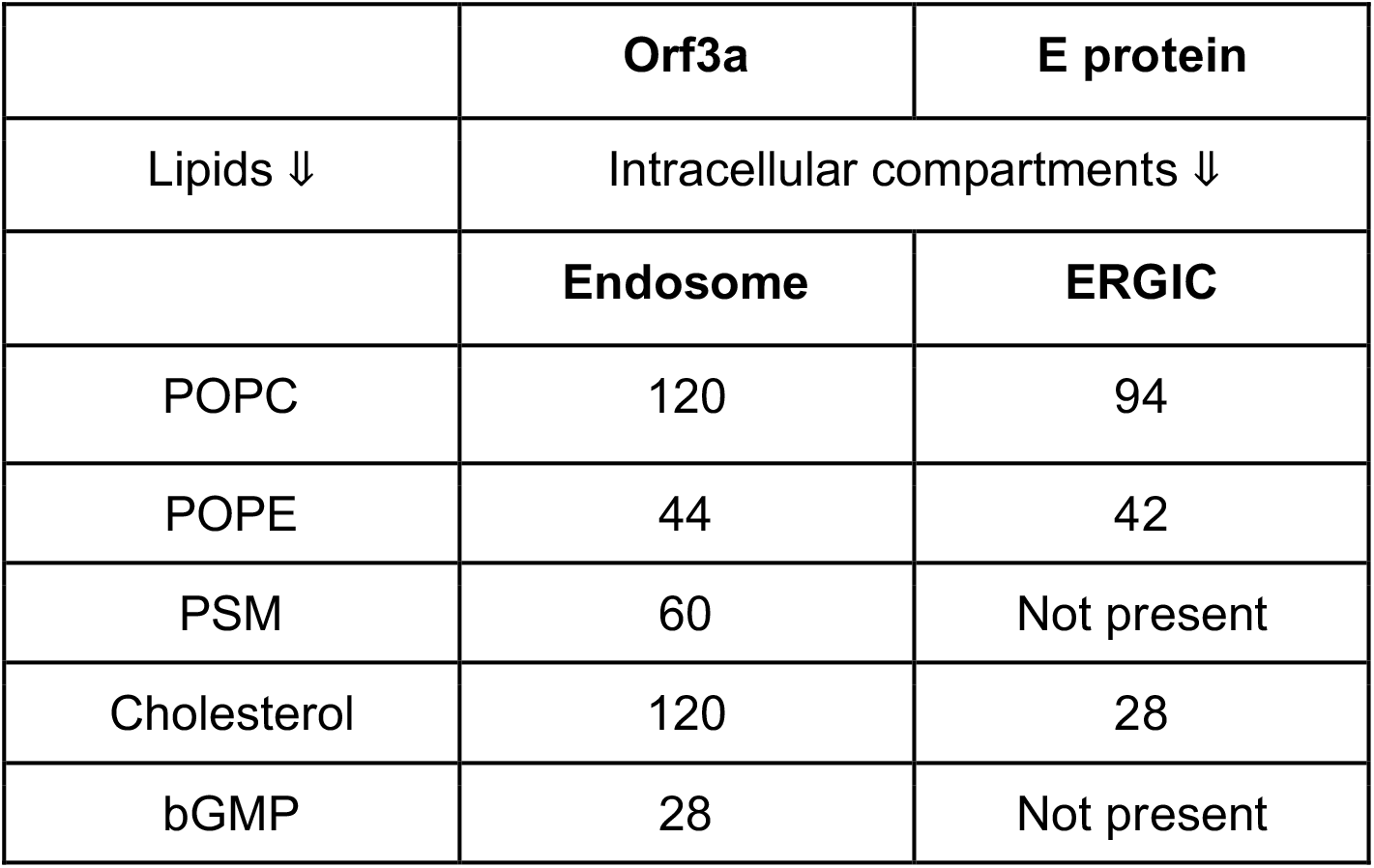
Lipid composition of different membrane components of intracellular compartments (endosome and ERGIC)

|  | **Orf3a** | **E protein** |
| --- | --- | --- |
| Lipids ↓ | Intracellular compartments ↓ |  |
|  | **Endosome** | **ERGIC** |
| POPC | 120 | 94 |
| POPE | 44 | 42 |
| PSM | 60 | Not present |
| Cholesterol | 120 | 28 |
| bGMP | 28 | Not present |

**Figure 3.**
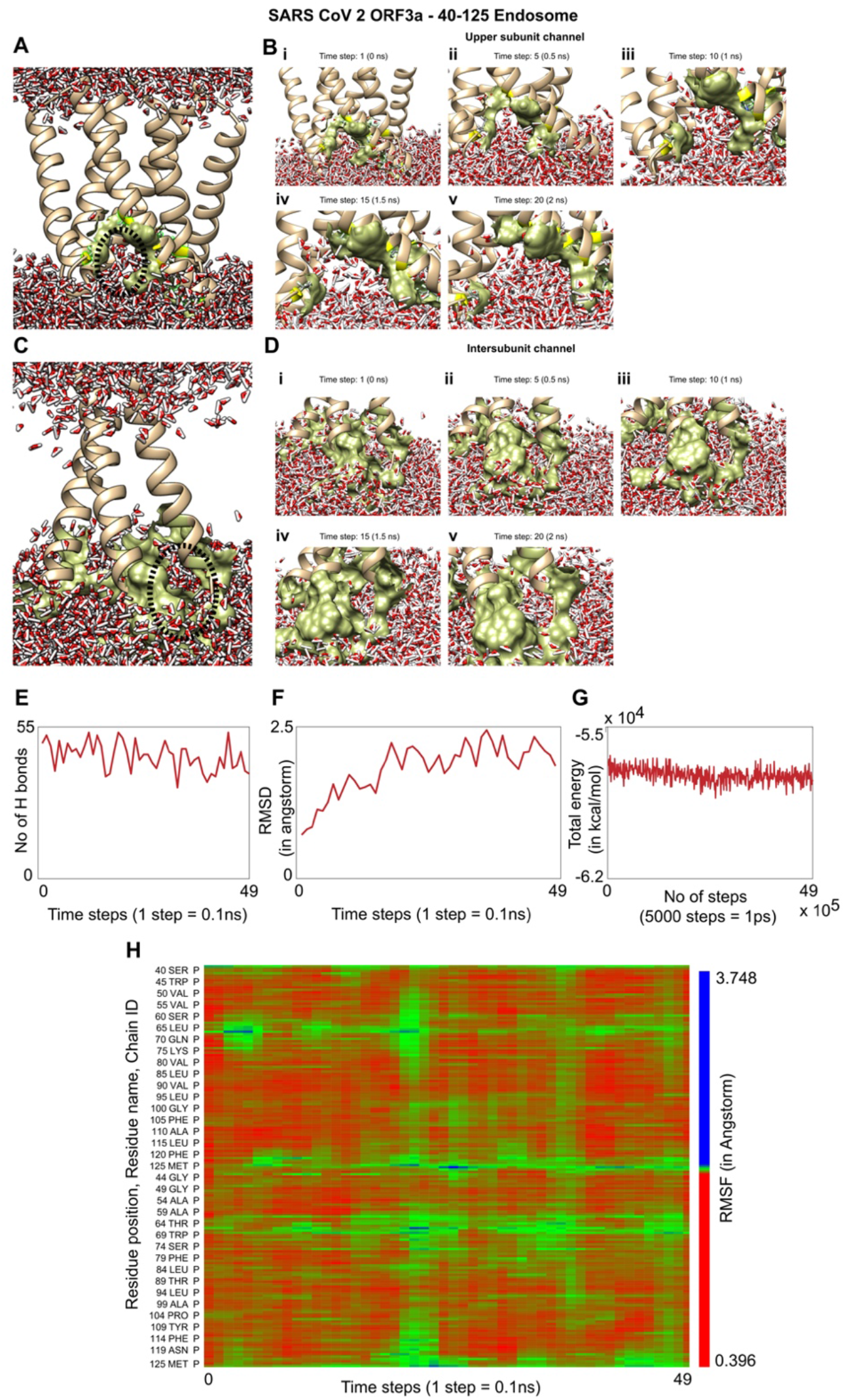
A. Insertion and equilibration of truncated Orf3a (40-125) from SARS-CoV-2 in a membrane system mimicking the endosomal compartment of a generalized human cell. The membrane is not shown in the figure to ease the visualization of the molecular machine. The upper channel has been pointed out using a dotted circle. B. Movement of water molecules through the upper channel at different time points (i) 0 ns (ii) 0.5 ns (iii) 1 ns (iv) 1.5 ns (v) 2 ns (Supplementary Movie 1) C. Insertion and equilibration of truncated Orf3a (40-125) from SARS-CoV-2 in a membrane system mimicking the endosomal compartment of a generalized human cell. The membrane is not shown in the figure to ease the visualization of the molecular machine. The lower channel has been pointed out using a dotted circle. D. Movement of water molecules through the inner subunit channel at different time points (i) 0 ns (ii) 0.5 ns (iii) 1 ns (iv) 1.5 ns (v) 2 ns (Supplementary Movie 1) E. Line plot showing the dynamics in the number of hydrogen bonds in the protein-membrane system as a function of time steps for 5ns where each time step is 0.1 ns. F. Line plot showing the RMSD of the protein-membrane complex and its change over a course of 5 ns where each time step is 0.1 ns. G. Line plot showing the total energy (kcal/mol) of the system in 5 ns of simulation with each 5000 steps=1 ps. H. Matrix representation showing the RMSF of individual residues of the protein over the time span of 5 ns where each time step is 0.1 ns.

### 3.4. Molecular dynamics of the TM region of SARS-CoV-2 E protein in an ERGIC membrane mimic shows water movement

The pentameric E protein from the SARS-CoV-2 has been modelled taking its solid-state NMR structure as the template (PDB id: 7K3G) (26), whose viroporin activity through the single channel formed by its TM region (8-40) has been analysed in a previous study (13) (Figure 4A). The protein-membrane complex was equilibrated energetically followed by 5 ns of MD simulation to analyse the continuous water channel formation through the proposed pore (Supplementary Movie 2). The water dynamics was observed at several intervals for 2 ns from the initial timestep starting at 0 ns (Figure 4Bi), 0.5 ns (Figure 4Bii), 1 ns (Figure 4Biii), 1.1 ns (Figure 4Biv), 1.2 ns (Figure 4Bv), 1.3 ns (Figure 4Bvi), 1.4 ns (Figure 4Bvii), 1.5 ns (Figure 4Bviii), and 2 ns ((Figure 4Bix). We observed that from 1 ns to 1.5 ns, the water molecules reach the proposed F26 bottleneck region (13) inside the pore of the E protein (Figure 4Biii-viii). We calculated the number of H-bonds (Figure 4C), RMSD (Figure 4D), total energy (Figure 4E), and RMSF (Figure 4F) for 5 ns of the simulation. The RMSD remained well below 5 angstroms (Figure 4D), indicating that the protein-membrane system has high structural stability. In addition, the total energy of the system remains relatively unfluctuating around -3.2 ×10^4^ kcal/mol (Figure 4E) throughout the time period of the simulation.

**Figure 4.**
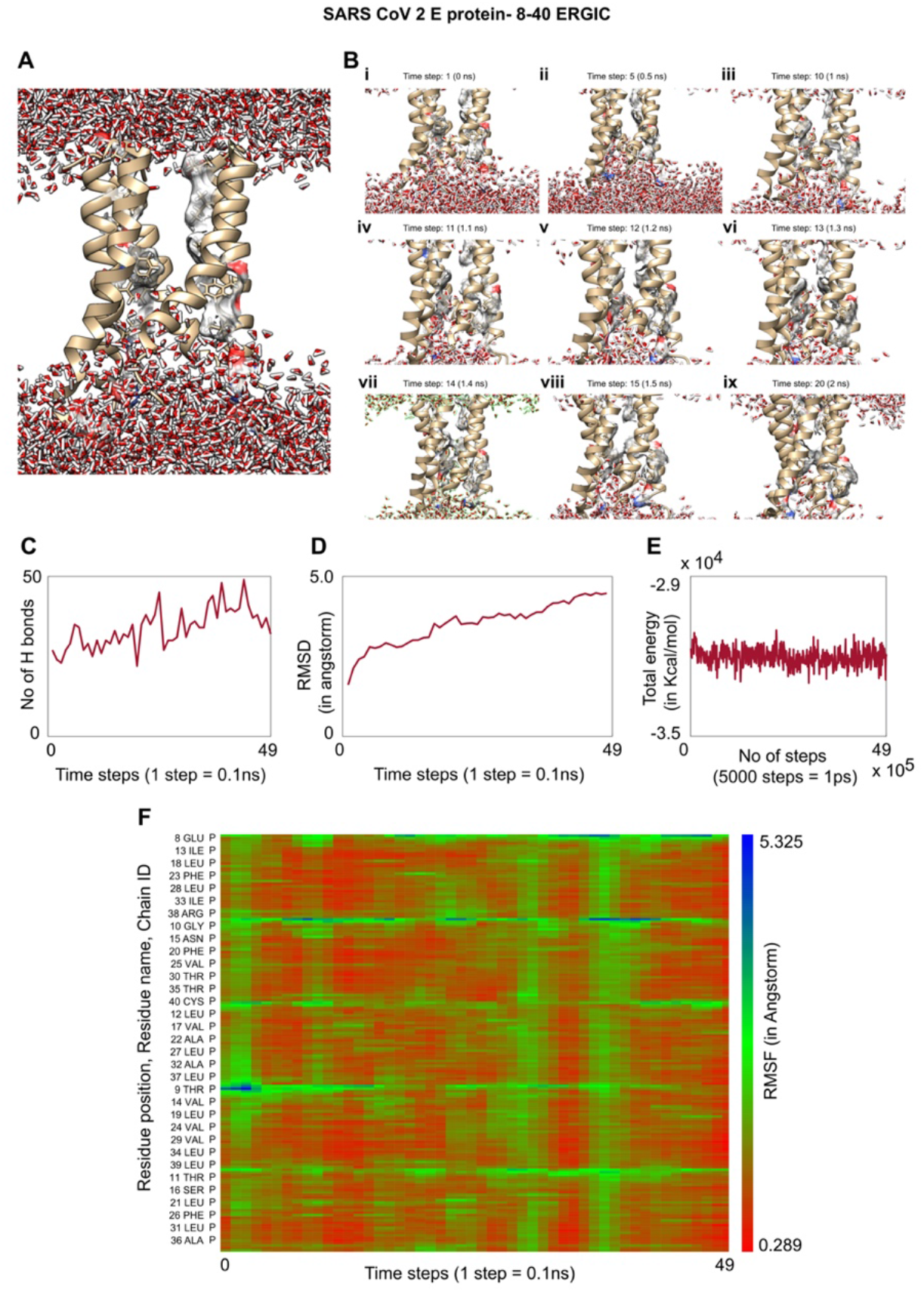
A. Insertion and equilibration of E protein (8-40) of SARS-CoV-2 in a membrane system mimicking the ERGIC of human cells. The membrane is not shown in the figure to ease the visualisation of the molecular machine. B. Movement of water molecules through the central pentameric channel of the TM region of the E protein at different time points (i) 0 ns (ii) 0.5 ns (iii) 1 ns (iv) 1.1 ns (v) 1.2 ns (vi) 1.3 ns (vii) 1.4 ns (viii)1.5 ns (ix) 2 ns (Supplementary Movie 3) C. Line plot showing the dynamics in the number of hydrogen bonds in the protein-membrane system as a function of time steps for 5ns where each time step is 0.1 ns. D. Line plot showing the RMSD of the protein-membrane complex and its change over a course of 5 ns where each time step is 0.1 ns. E. Line plot showing the total energy (kcal/mol) of the system in 5 ns of simulation with each 5000 steps=1 ps. F. Matrix representation showing the RMSF of individual residues of the protein over the time span of 5 ns where each time step is 0.1 ns.

### 3.5 Upregulated genes in COVID condition reflects cellular processes impacted by ion channelling activity

We used whole genome RNA sequencing data (36) to explore genes upregulated in SARS-CoV-2 infection, and whether they could be impacted by changes in cellular ionic homeostasis. PCA analysis (Figure 5A) suggested that uninfected controls and SARS-CoV-2 infected samples were mutually orthogonal to each other, suggesting that gene expression levels were indeed due to the infection. The first component explained 53% of the observed variance in the expression patterns. We further looked into the proportion of upregulated and downregulated genes (Fig 5B, 5C), and found 563 significantly upregulated genes and 75 significantly downregulated genes. The log of fold change (LFC) after the DESEQ2 pipeline was set at a minimum of 2, and adjusted p-value was set at 0.01. The top 50 upregulated and downregulated genes are shown in Supplementary Figure 2. PRKAA1 was upregulated in the infected samples, albeit it did not clear the threshold of LFC > 2 (Figure 5D) and interestingly, had been shown to regulate ion channelling activity of the host cell (37). We also observed significant upregulation in immunomodulatory genes like CD40, IFNL1, IFNL2, IFNL3, IL12A, IL33, IL6 and NFkB1 (Figure 5E). Among the downregulated genes were HSP90AB1, HSP90AA1 and HSP90B1 (Figure 5F). In our analysis, we focused on those genes only that could be impacted by or otherwise could impact cellular ion channelling and ionic concentration. In our analysis, 88% of the genes were upregulated while 12% were downregulated with our parameters (Figure 5G). In order to understand what cellular and molecular functions could be impacted by the upregulated genes, we constructed a gene network using Metascape (Figure 5H). The major functions implicated are defence response to virus, immune response regulating signalling pathway, response to IFN-β, regulation of cytokine production, response to IFN-γ, lymphocyte activation, SARS-CoV-2 innate immunity evasion, and regulation of I-κB-kinase/NFκB signalling (Figure 5H). All these processes imply the active participation and activation of the host defence system in case of a viral infection. We also evaluated the functional significance of protein-protein interactions of the upregulated genes and found that the following functions were enriched: defence response to virus, IFN-α/β signalling, cytokine signalling, post translational protein phosphorylation, calcium signalling pathway-Gα(q) signalling, PI3K-Akt-mTOR signalling, exocytosis and complement cascades (Figure 5I). Indeed, a lot of these functionalities could be directly or indirectly affected by ionic imbalances in the cell. These observations provide strength to our hypothesis that ion channelling activity by viroporins is responsible for viral pathogenesis and host cell responses in SARS-CoV-2.

**Figure 5.**
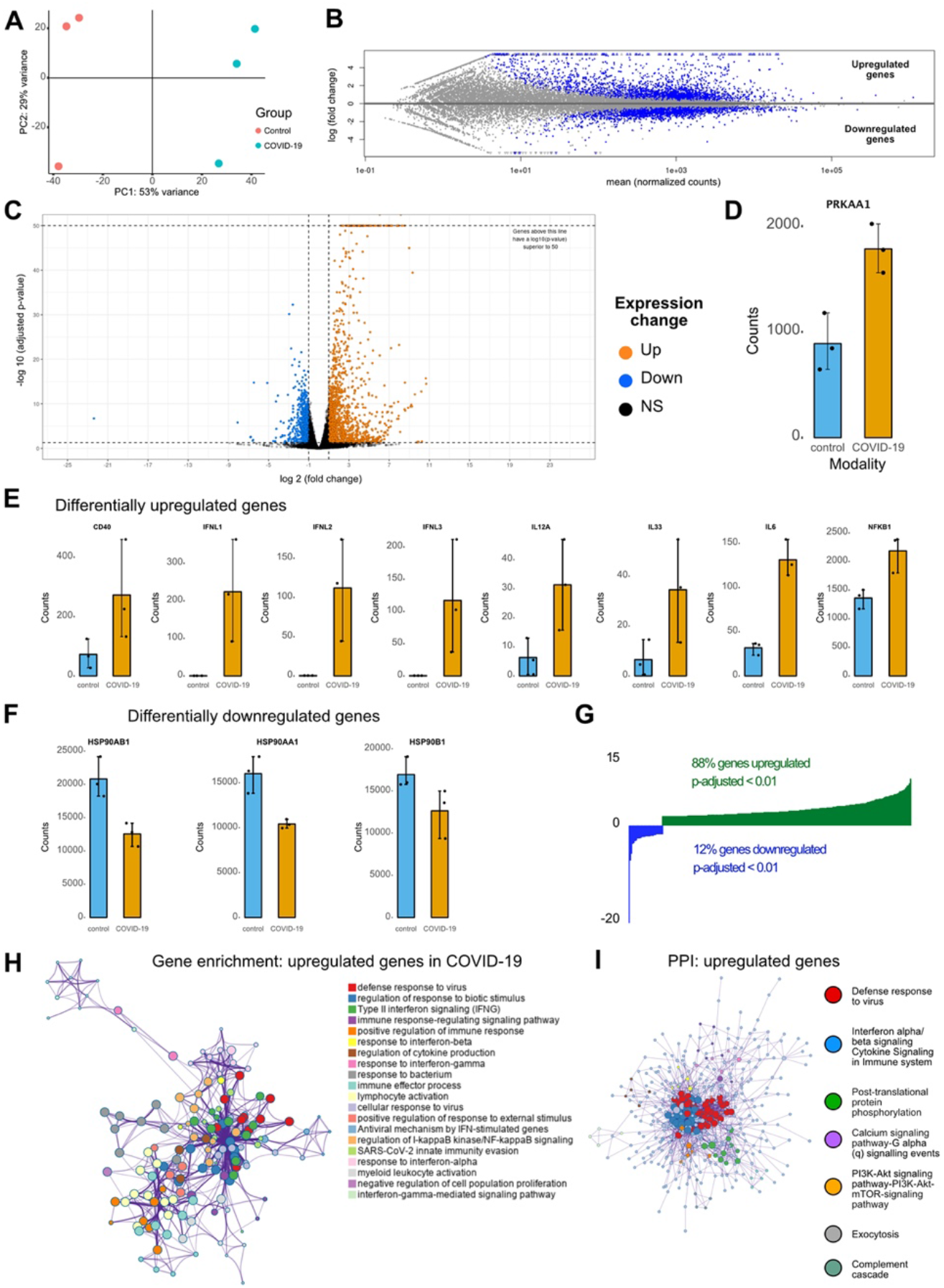
A. Principal component analysis (PCA) scatter plot representation of the variability in the dataset along the first two PC axes (PC1- 53% variability; PC2- 29% variability). Control and infected samples are orthogonal to each other. B. MA plot (M-log ratio of fold change; A- mean of normalized counts) for differentially significant gene expression in Infected samples compared to Controls. C. Volcano plot showing significantly upregulated genes (in orange), significantly downregulated genes (in blue) and non-significantly expressed genes (in black) between infected cells and controls. D. Bar plot showing gene counts of PRKAA1 gene between infected cells and controls. Error bars indicate Standard deviation. E. Example bar plots of differentially upregulated genes between infected cells and controls. The values plotted are gene counts with the error bars indicating standard deviation. Genes plotted: CD40, IFNL1, IFNL2, IFNL3, IL12A, IL33, IL6 and NFκB1. F. Same as E, but for differentially downregulated genes. Genes plotted: HSP90AB1, HSP90AA1 and HSP90B1. G. Distribution showing the percentage of genes significantly upregulated (p < 0.01) and with LFC >2. 88% of the genes were upregulated while 12% of the genes were downregulated. H. Gene enrichment analysis and gene ontology network obtained from upregulated genes in infected samples vs controls showing enrichment for immune functions and its associated signalling pathways. Metascape was used for this analysis. Different colored dots indicate different genes encoding the same function. I. Protein-protein interaction (PPI) enrichment analysis showing cellular functions determined by the upregulated genes in infected samples vs controls. Metascape was used for this analysis. Different colored dots indicate different genes encoding similar protein-protein interactions.

## 4. Discussion

Ion channelling activity is an important cellular event taking place in all organisms, from unicellular prokaryotes to multicellular eukaryotes like humans. Human cellular organisation and mechanisms of varied physiological events are directly or indirectly influenced by ion channels, which are mostly specific for the type of ion it transports. Almost all families of viruses encode one or more ion channel proteins which integrate in the host membrane and regulate key viral life cycle events like virion maturation, assembly and release. Viroporins oligomerize in the host membranes, leading to formation of permeable hydrophilic pores (23,38), which alters cellular ionic homeostasis in hosts. It leads to membrane depolarization and disruption of organelle architecture *via* membrane remodelling events, alteration of Ca^2+^ homeostasis (39) and protein trafficking.

SARS-CoV-2 has acquired several mutation hotspots in the Spike (S) protein across all its variants (40,41) which contributes to the increased pathogenicity of the variants (17). However, there are several other important viral proteins which play decisive roles in the viral life cycle and have remained conserved across the variants. We performed large scale sequence data analysis for the ion channelling viral proteins of SARS-CoV-2: Orf3a and Envelope protein, where we calculated the position specific frequency of each amino acid from sequences reported in the NCBI datasets for these two proteins. The positions harbouring the mutations reported from the analysis, were either identical to the residue in the corresponding position of the reference sequence with lower frequency, or different residue with higher frequency. These changes are random mutations and cannot be concluded as significant, as the sample space of the number of sequences were quite low in some of the variants. The variants with the highest number of sequences (Alpha and BA lineage of Omicron - Figure 1B) did not show any major mutations in the proteins. Nevertheless, we aligned the sequences of the variants and specified the random mutations in the ion channel forming region of the proteins. All the mutant structures were structurally aligned with the reference structure (PDB id: 7KJR) which showed no significant structural variability in the channel forming region. Afterwards, we looked into the membrane permeation and channel-forming mechanism of the Orf3a and E protein from SARS-CoV-2.

One of the most studied viroporins is the M2 channel of the influenza virus which is essential for viral replication and homeostasis. The M2 channel allows K^+^ ion influx and disrupts Na^+^/K^+^ ionic homeostasis in the late endosomes. It also acts as a proton channel in the TGN (pH∼6) which affects downstream protein trafficking machinery (42). The NS4A channel of HCV localises on the mitochondrial membrane and disrupts mitochondrial architecture by causing ionic imbalance in the organelle lumen (43). Viroporins activate apoptosis through the mitochondrial pathway via formation of apoptosome with pro-caspase 9 and apoptosis protease-activating factor-1 (44). P7 viroporin of HCV is a gated proton channel which causes H^+^ efflux, resulting in IL-1β production (45). The E protein of SARS-CoV-2 has been reported to rescue growth of K^+^-uptake deficient bacteria thus supporting its K^+^ conductivity. Additionally, it acts as a proton channel and causes bacterial cell death due to increased membrane permeabilization (28).

However, the exact molecular mechanism of these ion channelling events needs further exploration from a structural point of view. Our results demonstrate salient structural features which might determine how the viroporin functions. Hydrophilic pore formation is one of the fundamental features of a viroporin (23). We show formation of hydrophilic pore *via* water channel formation which could imply formation of ionic transfer mediums across the membranes. Such a passage medium through the upper and lower channels of dimeric Orf3a of SARS-CoV-2 is observed in human endosomal mimicking membranes. Endosome is an important structural organelle involved in a wide range of cellular functions. Water chain formation through the central pentameric pore of the Envelope (E) protein up to the proposed bottleneck region, defined the hydrophilic pathways through this viroporin of SARS-CoV-2 in human ERGIC membrane mimic.

The question remains, what impact do these viroporins impart at a physiological level? Indeed, our studies are *in silico* and have limitations of being non-experimental from an *in vitro* and *in vivo* standpoint. However, the impact of ionic imbalances in cellular micro-environment as a result of viral infections and viroporins have been studied in great detail earlier. One immediate observation comes from the previous SARS strain, the SARS-CoV-1 which showed that E protein localised in the ERGIC membrane, and facilitated the movement of Ca^2+^ ions into the cytosol (24). On the other hand, the Orf3a localised at the Endosome, Golgi apparatus and the plasma membrane transport K^+^ ions (46). A tight regulation of cationic and anionic ion channels controls the ionic homeostasis in the airways which can be correlated to complex pathological features in lung diseases (47). Viroporins localised in the subcellular membranes of these lung airway epithelial cells is the primary cause of ionic imbalance and thus can be potential therapeutic targets against ARDS, which is the primary reason for fatality in SARS-CoV-2 infection (48,49). Transcriptomics analysis from SARS-CoV-2 infected samples showed an upregulation in inflammatory response mediated by several interleukins and interferons (50) which are probably regulated by NFкB (51). Increase in CD40 (52), IL-6 (53), IL-12 (54) and IL-33 (55) transcripts strongly correlate with similar expression patterns of differentially expressed genes in Acute Lung Injury, ARDS and pulmonary fibrosis. NF-kB signalling has also been shown to be activated and induce inflammatory cytokines and chemokines, including IL-1b, IL-18, and IL-8 (56–58). AMPK is a master regulator of a wide spectrum of ion channels, carrier proteins and symporter-pumps (37), differential expression of which can impact their stimulatory and inhibitory effects (59). These channels, carrier proteins and/or pumps mediate the primary host cell response against viroporin action. Downregulation of ion channels is also known to be impacted by their interactions with Hsp proteins such as Kv7.4, a voltage gated potassium channel (60). Indeed, Hsp90, being a part of the protein folding machinery showed lower expression levels, probably due to disruption of the ERGIC and endosomal compartment. This is in line with our initial membrane disruption hypothesis as a result of change in ionic balance of the cell. In accordance with our study, E protein-mediated Ca^2+^ and K^+^ leakage, and Orf3-mediated K^+^ efflux could be a vital factor causing cellular ionic imbalance activating the NLRP3 inflammasome since, the activation of NLRP3 inflammasome directly correlates with the observed viroporin activity (46,61). Ionic imbalance in cells can promote build-up of reactive oxygen species (ROS) in the mitochondria, providing indirect to activation of NLRP3 (46). Our multivariate study on SARS-CoV-2 viroporins gives valuable structural insights into their mechanism of actions. We have elucidated the importance of Orf3a and E protein in COVID-19 pathogenesis on a structure-function association, potentially translating to changes in the transcriptome of the host cell. Further investigation along these lines can reveal potential therapeutic strategies against SARS-CoV-2.

## Supplementary materials

***Supplementary Figure 1: supplement to Figure 2***

Superimposition of experimentally determined Orf3a structure (7KJR, in orange) with the corresponding homology model structures of Orf3a (in blue) for the variants: (A) alpha, (B) Q lineage, (C) beta, (D) gamma, (E) delta, (F) AY lineage, (G) omicron and (H) BA lineage.

***Supplementary Figure 2: supplement to Figure 5***

Top 50 significantly upregulated (dark green) and downregulated (dark red) which have adjusted p-value of < 0.01 and LFC > 2

***Supplementary Movie 1:***

Molecular dynamics of TM- region (40-125) of Orf3a from SARS-CoV-2 inserted into a late endosomal-lysosomal membrane mimic.

***Supplementary Movie 2:***

Molecular dynamics of TM- region (8-40) of E protein from SARS-CoV-2 inserted into an ERGIC membrane mimic. The template of the modelled E protein is its NMR structure from SARS-CoV-2 (PDB id:7K3G).

## Authorship contribution statement

Manish Sarkar & Soham Saha (co-correspondence): Study design, structural modelling, molecular dynamics and analysis, bioinformatics. MS and SS drafted the manuscript and made the final figures.

Victor Hannothiaux: Data curation, big data analysis, residue frequency calculations, coding.

Paul Etheimer: Designed analysis automation for whole genome metagenome analysis using DeSEQ2 pipeline and performed the analysis on the transcriptomic data.

## Institutional Review Board Statement

**N/A**

## Informed Consent Statement

**N/A**

## Data Availability Statement

All our data are available upon request to corresponding authors

