## Supplementary figures and images for "SARS- CoV-2 viroporins: A multi-omics insight from nucleotides to amino acids"

### Supplementary Figure 1

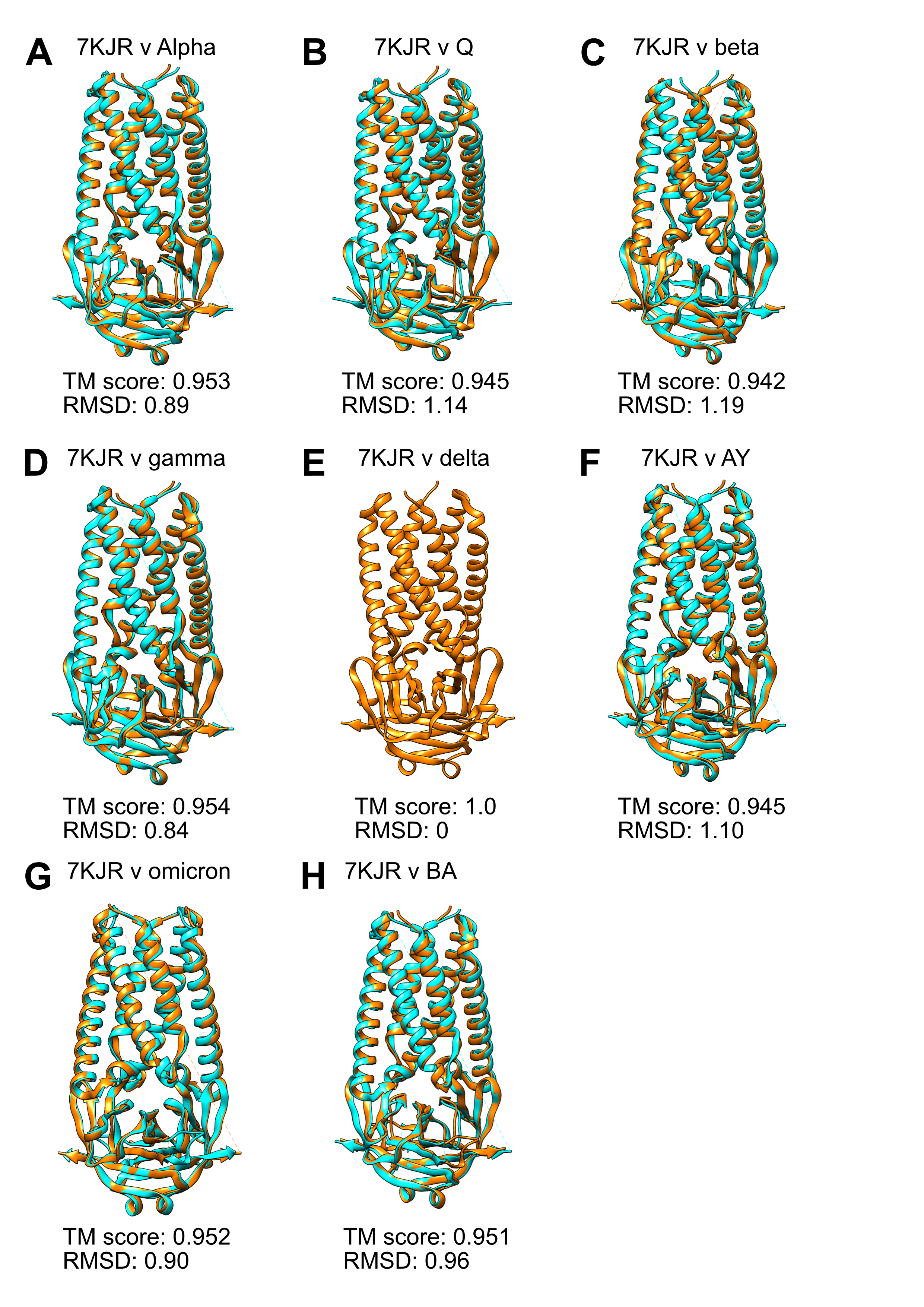

### Supplementary Figure 2

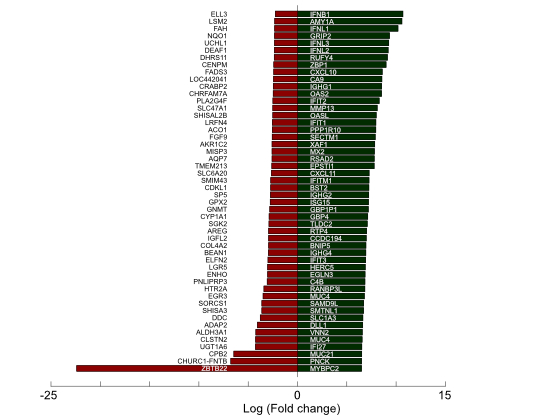
